# Interactions and collider bias in case-only gene-environment data

**DOI:** 10.1101/124560

**Authors:** Félix Balazard, Sophie Le Fur, the Isis-Diab collaborative group, Pierre Bougnères, Alain-Jacques Valleron

## Abstract

**Background:** Case-only design for gene-environment interaction (CODGEI) relies on the rare disease assumption. A negative association due to collider bias appears between gene and environment when this assumption is not respected. Genetic risk estimation can quantify part of the predisposition of an individual to a disease.

**Methods:** We introduce Disease As Collider (DAC), a new case-only methodology to discover environmental factors using genetic risk estimation: a negative correlation between genetic risk and environment in cases provides a signature of a genuine environmental risk marker. Simulation of disease occurrence in a source population allows to estimate the statistical power of DAC and the influence of collider bias in CODGEI. We illustrate DAC in 831 type 1 diabetes (T1D) patients. Results: The power of DAC increases with sample size, prevalence and accuracy of genetic risk estimation. For a prevalence of 1% and a realistic genetic risk estimation, power of 80% is reached for a sample size under 3000. Collider bias offers an alternative interpretation to the results of CODGEI in a published study on breast cancer.

**Conclusion:** DAC could provide a new line of evidence for discovering which environmental factors play a role in complex diseases or confirming results obtained in case-control studies. We discuss the circumstances needed for DAC to participate in the dissection of environmental determinants of disease. We provide guidance on the use of CODGEI regarding the rare disease assumption.

Key messages
- A complex disease is a collider between genetic determinants and environmental factors; it is a consequence of both.
- Collider bias can affect the results of the case-only design for gene-environment interactions when the disease is common.
- Using genetic risk estimation, collider bias can be used to discover or confirm the association between a disease and an environmental factor in a case-only setting.
- Statistical power of this approach is small when the disease is rare. Power increases with sample size, prevalence and genetic risk prediction accuracy.

## Introduction

Case-only gene-environment studies are attractive since data are often easily available in cases. It also means that the selection of controls, a sensitive process, can be avoided. Case-only design for gene-environment interaction (CODGEI) allows to study gene-environment interactions in this setting^1,2^. Here, we propose DAC, a new methodology that uses the same data. The two methodologies are complementary as DAC has statistical power when the rare disease assumption of CODGEI is violated.

CODGEI uses case-only data to identify gene-environment interaction. Specifically, if both G and E are binary traits as shown in Table 1, the cross-product ratio (CPR) ad/bc computed from the case-only data is an estimator of the interaction odds-ratio OR_I_ between G and E.

**Table 1:** Gene-environment data in a case-only setting.

|  | G=0 | G=1 |
| --- | --- | --- |
| E=0 | a | b |
| E=1 | c | d |

CODGEI needs two assumptions to be applied: G and E must be independent in the general population^3,4^ and the disease must be rare at all levels of gene and environment: for all g and e, P(D=1|G=g,E=e)<<1

When this second assumption is not valid, collider bias appears between gene and environment and may jeopardize the interpretation of studies applying CODGEI.

Collider bias (or collider-stratification bias) is the negative correlation that appears between two causes when conditioning on their shared consequence (the collider)^5^. It can mislead epidemiological investigation^6,7^. A classic example is Berkson’s bias in which two diseases are negatively associated in a hospitalized population even though they are independent in the general population^8,9^. In this example, the collider is hospitalization, the shared consequence of both diseases. By looking only at patients in the hospital, i.e. by conditioning on hospitalization, a negative correlation appears between the two diseases.

In the setting of case-only gene-environment data, if both the gene and the environmental factor under consideration are causes of the disease, then conditioning on the disease, i.e. considering only cases, a negative association appears between gene and environment. This principle is illustrated in Figure 1a and 1b. However, it is not necessary for the environmental factor to be a cause for collider bias to appear. If the environmental factor of interest is simply correlated with a causal factor for the disease, collider bias will appear as shown in Figure 1c and 1d. No causal claim can therefore be made.

**Figure 1:**
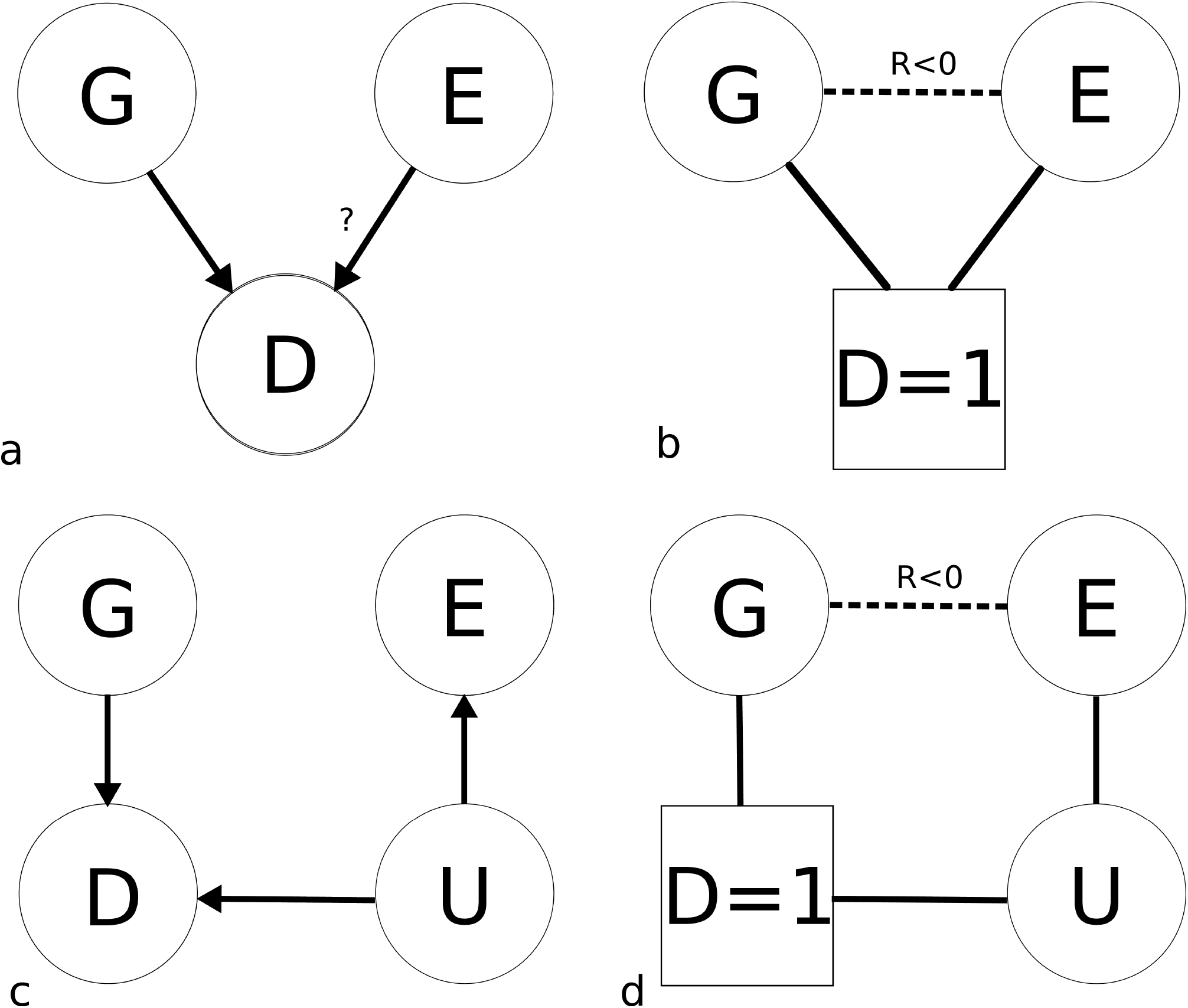
Collider bias in case-only gene-environment data, a: In the general population, disease is a consequence of both genetic and environmental causes. Depending on the environmental factor considered, we can assume independence between gene and environment. b: When considering only cases ie conditioning on the disease, a negative association appears between genetic risk and the environmental factor. c and d: If the environmental factor E is only a marker for the unobserved U that is the true environmental cause, collider bias appears nevertheless between G and E.

However, Piegorsch et al.^1^ showed is that when the rare disease assumption is verified, the CPR estimates the interaction odds-ratio OR_I_ and therefore collider bias is not present. The impact of deviations from the rare disease assumption has been studied by Schmidt and Schaid^10^. These authors noted that the CPR’s asymptotic value is the interaction risk ratio RR_I_. The RR_I_ measures the departure from multiplicative risk ratios. However, the interaction term that is estimated by a case-control study is an interaction odd-ratio OR_I_ that measures the departure from multiplicative odd-ratios, i.e. an interaction in the logistic model. Under the rare disease assumption, RR_I_=OR_I_. Schmidt and Schaid evaluated the influence of deviations from the rare disease assumption on the mismatch between OR_I_ and RR_I_. Their conclusion was that RR_I_ can be substantially smaller than OR_I_ under large deviations from the assumption. *We believe that this conclusion is somewhat misleading.* Since ratios are on a multiplicative scale, an underestimation would mean that RR_I_ is closer to 1 compared to OR_I_. In their Figure 3, you can see that when OR_I_=1, we have RR_1_<1. This means that when there is no interaction in the logistic model, an inverse interaction will be detected by CODGEI, in other words a negative association appears between G and E. This negative association is expected when one considers collider bias.

CODGEI estimates correctly under all circumstances RR_I_. However, meta-analysis need to collate estimators of the same quantity. Having RR_I_ for case-only studies and OR_I_ for case-control studies is problematic as the null hypotheses of no interaction are different in the two quantities. To document an example where collider bias is the driving force hidden behind epidemiological results, we searched in the literature for a study that applied CODGEI in a situation where the rare disease assumption is not respected. The study that best suited our criterion is the Genetic and Environmental Modifiers of BRCA1/BRCA2 Study (GEMS)^11^. It deviates strongly from the rare disease assumption as it considers interactions between the highly penetrant BRCA1/2 and environment in breast cancer, the most common cancer in women. We show below that the conclusions of the study are changed if we define the null hypothesis as multiplicative odd-ratios and not multiplicative risk ratios.

In the example of CODGEI and also more generally in epidemiology, collider bias is seen as a nuisance that hinders understanding. Herein we propose a change of view-point in order to harness collider bias in service of epidemiology. In order to do this, we need to maximize collider bias and therefore deviate as much as possible from the rare disease assumption. This can be achieved if, instead of considering one variant at a time, we consider genetic risk predictions that uses many variants to estimate as accurately as possible the genetic risk. Indeed the individuals with the worst combination of variants will have a non-negligible risk of disease. Genome-Wide Association studies (GWAS) datasets have been used to estimate genetic risk using statistical learning techniques^12-17^. Genetic risk predictions have been evaluated for T1D, Crohn’s disease and celiac disease. For those three diseases, similar prediction accuracy was achieved with area under the receiver operating curve (AUC) around 0.85^12,13,15^.

By conditioning on disease, i.e. by considering only cases, a negative association found between the genetic risk and an environmental candidate will signal a true association between the environmental marker and the disease. We refer to this methodology as Disease As Collider (DAC). To sum up, DAC allows to detect or confirm a putative environmental marker by looking for an association of this factor with genetic risk in case-only data.

In the next section, we describe the method and its assumptions. We illustrate our methodology on a subset of genotyped patients from the Isis-Diab case-control study of T1D focused on the search of environmental factors^18^. Then, we present a simulation framework of disease occurrence as a function of the individual genetic susceptibility and environmental risk that allows to estimate the power of DAC as well as to evaluate the influence of collider bias in CODGEI. Using this framework and relying on the genetic risk distribution from our illustration data, we estimate power for our illustration and in generic scenarii. We evaluate the influence on power of prevalence, prediction accuracy of the genetic risk estimation and sample size. Using the same simulation framework, we also show the influence of collider bias in the GEM Study that applied CODGEI on breast cancer^11^.

## Methods

### Disease as collider

#### Model assumptions

##### Genetic risk and environmental marker independence

In order to attribute to collider bias the responsibility for an association between genetic risk and an environmental factor in cases, we need to assume that genetic risk and the environmental factor are independent in the general population. This assumption is shared with CODGEI.

##### Multiplicative combination of odd ratios

We need to make an assumption on how genetic risk and environmental risk combine in order to evaluate the power of DAC. In order to express our assumption, we need a few definitions. The logistic model transfers probabilities in [0,1] to log odd-ratios in the real line thanks to the logit function logit(x)=log(x/(1-x)). We refer to the target set of the logit function as the logit scale. For an individual with genome G (e.g. measured through genotyping) and environment E (e.g. measured through responses to an environmental questionnaire), we can define its genetic risk of developing a disease D by R_g_(G)=logit(P(D=1|G)) and its environmental risk R_e_(E)=logit(P(D=1| E)). R_g_ and R_e_ are actually the logit of an absolute risk. As most of our calculations are made on the logit scale, we will write risk instead of risk on the logit scale throughout.

The simplest way to define the total risk R(G,E)=logit(P(disease|G,E)) is to assume that environmental and genetic odd ratios combine multiplicatively, i.e. environmental and genetic risk combine linearly on the logit scale. We therefore have that:

R(G,E)=logit(P(disease|G,E))=R_g_(G)+R_e_(E).

This is an assumption of absence of interaction between the genetic risk and the environmental risk. It is the main difference between DAC and CODGEI as the latter attempts to detect such interactions.

#### Description of DAC

Under the assumptions of independence between gene and environment and multiplicativity of odds-ratio, there is an association between genetic risk and environmental factors in cases due to collider bias. Our method consists simply in estimating the genetic risk in cases and then on testing for association between genetic risk and environmental factors using standard tests (such as a linear regression t-test) while controlling for potential confounders.

The association that appears because of collider bias is a negative association. Therefore, DAC predicts that the cases the most at risk genetically are the least at risk because of environment. When a putative direction of association has been established, one can perform one-sided tests. This is the case when DAC is applied to confirm findings from a case-control association.

#### Application of DAC to the Isis-Diab study

The Isis-Diab cohort is a multi-centric study of T1D patients in France which recruitment started in 2007. We used a genetic score presented in the article by Wei et al^12^ and trained on the WTCCC1 data^19^. Using this genetic risk score, we applied DAC on 7 questions on the environment before diagnosis which had been associated with T1D in the previously published case-control study^18^. 831 patients were used in the analysis. Detailed description of the data, the analysis and the results are in the supplementary material.

### Framework for estimating influence of collider bias in a case-only study

Quantifying collider bias in a case-only study allows us to estimate the power of DAC and also to assess the bias in the application of CODGEI. The assumptions made for DAC correspond to the null hypothesis of no interaction on the logit scale that CODGEI tests. In both cases, we simulate disease occurrence in a source population according to our model assumptions. For a sample size of N patients, the source population consists of N/K individuals, K being the prevalence of the disease. To allocate disease status in this synthetic population, we need to define a genetic risk distribution and an environmental risk distribution. We describe the choice of distributions used for the different tasks below. Once both distributions are defined, we then attribute to each individual in the population its genetic risk and its environmental risk by drawing independently from those distributions. This uses the assumption of independence of genetic risk and environmental factor in the population.

Once both a genetic risk and an environmental risk are defined for each individual in our source population, we define the total risk as the sum of the two risks in accordance with our assumption of multiplicative combination of odd-ratios.

To decide whether an individual with genes G and environment E has the disease, we draw a uniform variable U on [0,1] and we could then define the disease variable D:

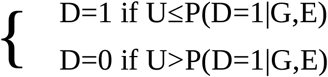

This approach would yield a different number of cases in each simulation. To always have N patients, we compute R(G,E)-logit(U) and define the top N individuals for that sum as the patients (D=1). The distribution of logit(U) where U is uniformly distributed over [0,1] is called the Laplace distribution.

Finally, when the simulated sample has been defined, we compute from it the quantity of interest and store it. We then repeat the procedure the desired number of times. We then have a distribution of the quantity of interest.

In the case of estimation of power of DAC, on each simulated sample, we perform a regression of the environmental factor on the genetic risk in the patients and obtain our quantity of interest: a onesided p-value. Our estimator of power is then the proportion of p-values under the threshold 0.05.

In the case of the influence of collider bias in CODGEI, we compute the cross-product ratio (CPR) on each simulated sample. With the resulting empirical distribution of CPR under the null hypothesis of no interaction on the logit scale, we can define a rejection region as the complementary of the 95% confidence interval of the CPR under the null.

We now turn to the definition of the genetic and environmental risk distribution and the choice of parameters in each setting.

#### Genetic risk distribution

The distribution of genetic risk in the general population is a mixture of the distribution of genetic risk in the controls and in the patients. If we denote D(X) the distribution of X, we have that:

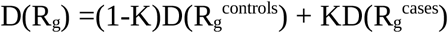

It should be noted that an individual whose genetic risk comes from the distribution of genetic risk for cases does not necessarily have the disease. In practice, we used the distributions of genetic risk obtained in our application of DAC on the Isis-Diab study (cf supplementary material and supplementary figure 1). The genetic risk estimation has been calibrated in order to represent probabilities. We sample N genetic risks from the genetic risks of Isis-Diab patients and we sample the rest from the genetic risks of the controls (cases of non-auto immune diseases in the WTCCC1 data).

In the case of a single variant such as BRCA1 or BRCA2, the distribution in the general population is given by the prevalence of the mutations. Its risk on the logit scale can be obtained from the prevalence of breast cancer in the general population and in carriers of the mutation. We chose a prevalence of breast cancer of 12 %, a prevalence of breast cancer among carriers of BRCA1/2 of 60%^20^ and therefore the OR for BRCA was 11. We chose a prevalence of 0.1 % for BRCA1 and 0.2 % for BRCA2^21^.

#### Power estimation in generic scenarii

For the estimation of power in the generic scenarii, we evaluated the influence of prevalence and also of prediction accuracy of genetic risk. To do this for prevalence, genetic risk was left untouched, we set prevalence to 0.2%, 0.6% or 1% and sample size to 500, 1500, 3000 or 5000. The three prevalences correspond to the prevalence of T1D in France for the lowest, T1D in Finland for the intermediate value and high estimation of prevalence of celiac disease for the highest^22^. Concerning the influence of prediction accuracy of the genetic risk, we set prevalence at 0.2%, we modified the genetic risk estimate to have an AUC of 0.88, 0.90 or 0.92 and we set the sample size to 500, 1500, 3000 or 5000. The genetic risk distribution with modified AUC was obtained by adding to the risk of patients a constant chosen to obtain the desired AUC. The estimate of risk in patients and controls was then calibrated again to correspond to probabilities.

Concerning the definition of the environmental risk, we chose an effect size of 3 which is a large but plausible effect size for epidemiology and we chose the most favorable distribution of the environmental factor in the patients, i.e the one with the most variance: an evenly split binary variable. The distribution in the source population was weighted by the inverse of the relative risk to obtain the desired distribution in cases.

#### Estimating collider bias in GEMS

We investigated if the four significant associations reported in the GEM study could be explained by collider bias. The four associations were BRCA1 and alcohol use (yes vs no), BRCA1 and parity (nulliparous vs 3 children or more), BRCA2 and parity (nulliparous vs 2 children) and BRCA2 and age at menarche (before 11 vs after 14). Given the main effects in the literature for the three risk factors, the RR_I_ were in the direction expected from collider bias. We used the sample size and the number of carriers relevant for the 4 comparisons. We chose a RR of 1.32 for alcohol use vs no alcohol^23^, a RR of 1.29 for nulliparous vs 2 children, a RR of 1.54(=1.29/0.84) for nulliparous vs more than 3 children^24^ and a RR of 1.05^-4^ for menarche after 14 years old vs menarche before 11^25^. In the same fashion as above, the distribution of the risk factor in the source population was weighted by the inverse of the relative risk to obtain the observed distribution in cases.

The general procedure presented above had to be adapted to the precise setting of the GEM study. Indeed, all patients were not included in the study: they included all patients carrying BRCA1/2 and a number of non-carrier patients as a comparison group. In consequence, the prevalence of BRCA1/2 is much higher in the study than in the population of patients. To take this into account, we add an additional step to the simulation after the allocation of disease status. We define a variable for inclusion in the study. All patient carriers are included in the study and the rest of the sample size is filled at random from non-carrier patients. We then use only the patient who are included in the study. This means that the source population is larger than N/K. We adjust the size of the source population to obtain in average the observed fraction of carriers in the simulated samples.

#### Code

Code used for analysis and power estimation is available at github.com/FelBalazard/DAC.

## Results

### Power estimation

The results of the power estimation in generic scenarii are presented in figure 2. Power increases with sample size, prediction accuracy of the genetic risk and prevalence of the disease.

**Figure 2:**
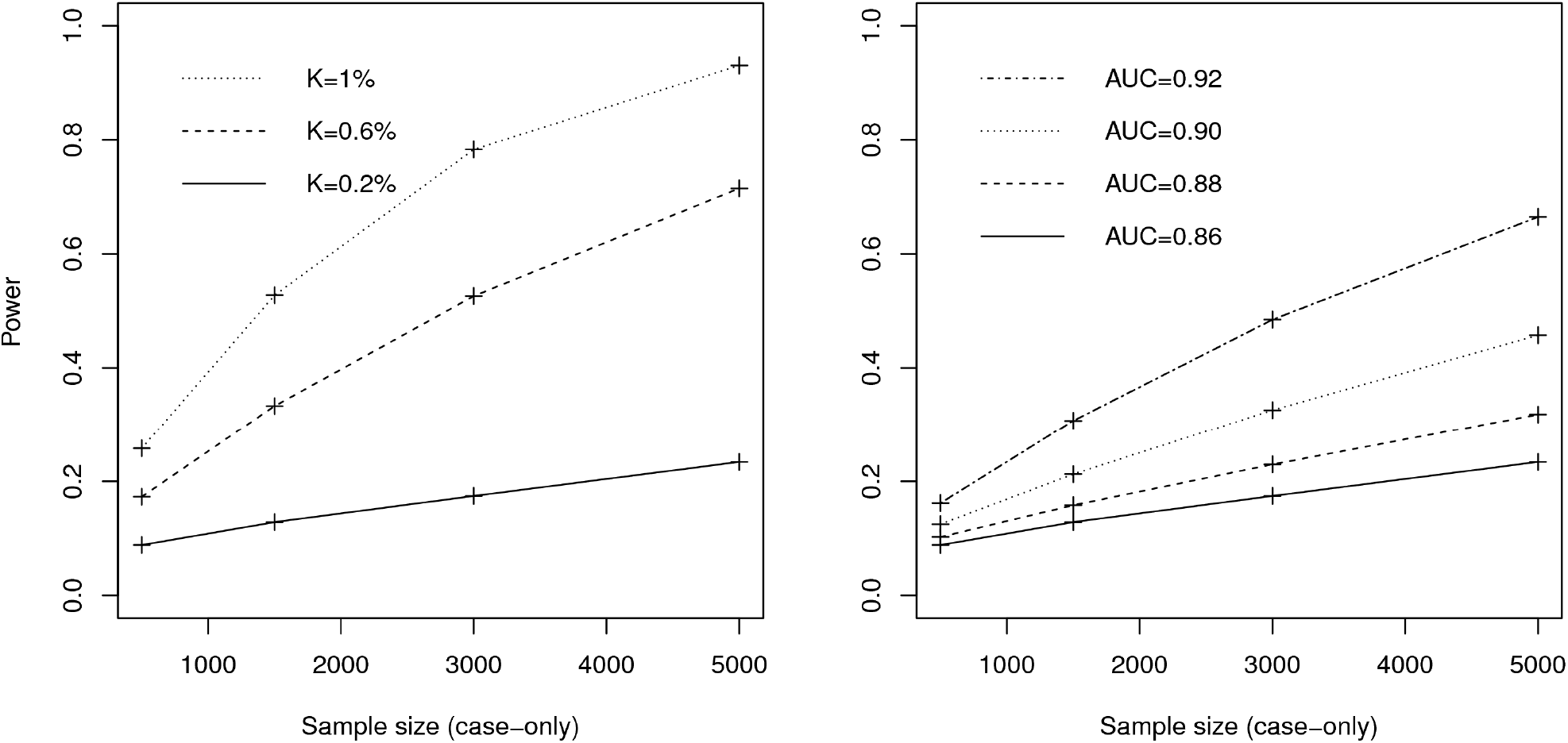
Power of the DAC methodology in different settings. The effect size is set at 3 and the environmental factor in cases is evenly split. a: Influence of the prevalence of the disease on power. The AUC of the genetic risk estimator remains at 0.86. b: Influence of the genetic risk accuracy (AUC) on power. The prevalence of the disease remains at 0.2%

With a prevalence of 0.2% and an AUC of 0.86, power was very limited. Even if our sample size had been 5000 cases and despite the favorable assumptions made on the effect size and the distribution of the environmental factor in cases, power would be only 26%.

Our estimation show that power depends strongly on prevalence of the disease. For a disease with prevalence of 1%, 80% power is attained for a sample size under 3000.

### Collider bias in the GEM Study

The results are presented in Table 2. The median and a 95% CI for the CPR under the null hypothesis of multiplicativity on the odds-ratio scale is presented alongside the CPR adjusted for age and center from the original article. In the 4 cases, there is a shift away from 1 of the median CPR and the reported CPR falls in the 95% CI. This means that the results from the GEM Study are not significant under the null hypothesis used in case-control studies.

**Table 2:**
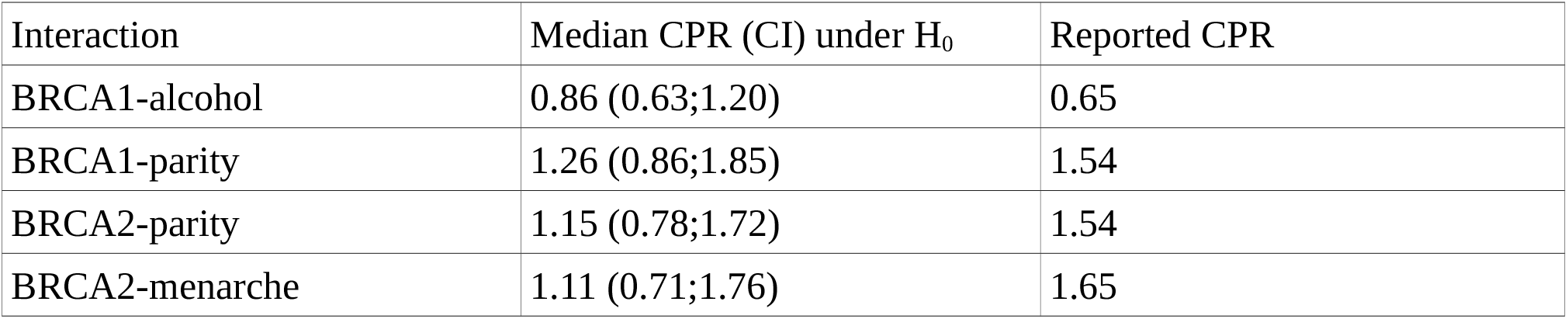
Results of simulations under the null hypothesis of no interaction on the logit scale for the 4 significant associations in the GEM Study. The reported CPR is reproduced from the original paper.

## Discussion

CODGEI has been proposed in 1994 to uncover gene-environment interaction using case-only data^1^. Here, we propose DAC as a new methodology for analysis of case-only data; it allows to discover or confirm associations of environmental factors with disease using genetic risk estimation. Ideally, DAC should be used after a standard environmental case-control study to confirm findings.

As important information is missing in the case-only setting, assumptions need to be made to be able to draw conclusions from case-only data. Both DAC and CODGEI rely on an assumption of independence between gene and environment. This is reasonable but deviation from independence should be kept in mind as an alternative explanation for a positive result. While certain genes affect certain exposures such as alcohol consumption^26^, coffee consumption^27^ or smoking^28^, there is a priori for independence between most genes and most environmental factors. We stress that the only independence needed for DAC is between the aggregated genetic risk score and the environmental factor: DAC does not require independence between each SNP and the environmental factor. When the environmental factor has genetic determinants and case-control data is available, Mendelian randomization^29^ will be more informative as it allows to substantiate causal claims.

Under the assumption of independence, two phenomenons are present in the case-only gene-environment data: interactions and collider bias. If there is no interaction between genetic risk and environment, only collider bias is left and DAC can be applied. This means that DAC is dependent on an assumption of absence of interaction. Indeed, interactions between genetic risk and environmental factor are problematic for DAC. A negative interaction strengthens the negative association that DAC tries to uncover but makes the findings less actionable as the people at highest genetic risk would respond less to intervention on the environmental factor (if the factor is a cause and not a mere marker). A positive interaction cancels the negative association that DAC tries to uncover despite increasing the prevention potential of the factor. This is a notable caveat to DAC as interactions between an aggregated genetic risk and environmental factors have been detected in relation to obesity^30^ and must be present in other settings as well.

Under the rare disease assumption, collider bias is negligible and therefore only interactions are left to explain an association between gene and environment in the case-only data. This is the rationale for CODGEI. As a consequence, genetics are considered differently in the two methods. For DAC to be successful, we need to maximize the distance to the rare disease assumption and therefore we use an aggregated risk estimation. On the other hand, in CODGEI, each variant is considered on its own.

This theoretical argument for absence of collider bias at low prevalences is in accordance with the results of power estimation. Those power estimations show that DAC can be successful in higher prevalence situations, with large sample sizes and better genetic risk estimation. However, in more common diseases, genetic risk estimation typically obtains sensibly weaker results and the prospective cohort design is more feasible. Nevertheless, DAC needs stronger prevalences of the disease to achieve reasonable power. This could be obtained in countries where T1D has a high prevalence such as Finland or on more frequent diseases such as celiac disease.

Given the prevalence of T1D in France, DAC is underpowered in the setting of the Isis-Diab study as shown in the supplementary material. Nevertheless, the application of our method to these data illustrates the practical considerations that go into applying DAC such as the problem of confounding by age at diagnosis. Furthermore, it allowed to base our power estimations on an actual predicted genetic risk distribution.

DAC underscores the importance for epidemiology of having a genetic risk estimation as predictive as possible. There has been limited access to the largest consortium datasets for this goal and a consequent turn to methods that use only summary statistics^16,31^. In the case of Crohn’s disease and ulcerative colitis, the International Inflammatory Bowel Disease Genetic Consortium dataset was used for this purpose and significant improvement was obtained^13^. Methodology to adapt machine-learning methods to GWAS datasets is also a promising avenue of research^17^.

CODGEI is quite resistant to the rare disease assumption. For example, interaction between BRCA1/2 and oral contraceptives in ovarian cancer^32^ are only marginally affected by collider bias despite the high penetrance of BRCA1/2 (results not shown). However, assuming one is interested in multiplicativity of odd-ratios and not the less used multiplicativity of relative risks, interactions between risk factors and BRCA1/2 in breast cancer in the GEM Study^11^ can be explained by collider bias. When the disease is common and there is highly penetrant variants, CODGEI should be applied with caution and collider bias should be considered as an alternative explanation for a significant negative association between gene and environment in cases. If the prevalence of the disease and the main effects of both genetic variant and environmental factor are known, it is possible to use the simulation framework that has been described here to test the presence of interaction on the logit scale.

## Ethics

The research protocol of the Isis-Diab study was approved by the Ethics committee of Ile de France (DC-2008-693) and the Commission Nationale Informatique et Libertés (DR-2010-0035). The ClinicalTrial.gov identifier was NCT02212522. All patients provided written informed consent for participation in the study and donation of samples.

## Acknowledgements

We thank Gérard Biau for his comments on the manuscript. We thank Mark Lathrop for the Isis Diab genetic data. We thank Yoichiro Kamatani for performing imputation on the Isis-Diab genetic data. We thank Sophie Valtat for rationalizing the questionnaires. We thank Alain Fourreau, Adeline Guégan, Gaël Leprun and Valérie Jauffret for technical assistance in the epidemiological investigation. We acknowledge the collective effort of the Isis diab collaborative group. We thank the participants and their parents for their time.

We thank the Wellcome Trust Case Control Consortium (WTCCC) for making the Affymetrix data available for our analysis. The WTCCC is funded by Wellcome Trust award 076113, and a full list of the investigators who contributed to the generation of the data are available from http://www.wtccc.org.uk.

